# Volumetric Functional Ultrasound Imaging in Macaques

**DOI:** 10.64898/2026.04.27.721041

**Authors:** Nora E. Fitzgerald, Gabriel Montaldo, Mathilda Froesel, Alan Urban, Wim Vanduffel

**Author notes:** These authors contributed equally (Nora E. Fitzgerald, Gabriel Montaldo).

## Abstract

Linking circuit level activity to large scale functional organization requires imaging methods combining high spatial resolution, broad coverage, and single trial sensitivity. We present volumetric functional ultrasound imaging (3D-fUS) in behaving macaques, enabling imaging of ~1 cm^3^ cortical volumes at high spatiotemporal resolution (100 × 150 × 150 μm^3^ voxels, 1.67 Hz). Visually evoked responses were reliably detected at the level of single trials and single voxels, substantially reducing experimental time. To enable model-based analyses analogous to functional magnetic resonance imaging (fMRI), we estimated a canonical fUS hemodynamic response function (fUS-HRF) that was consistent across subjects, cortical areas, and visual stimuli and was well approximated by a gamma function. Compared with canonical fMRI HRFs, the fUS-HRF exhibited faster dynamics, enabling shorter and more closely spaced stimulus presentations. Together, these results establish 3D-fUS as a fast, volumetric, and circuit relevant imaging modality for efficient investigation of distributed cortical dynamics in primates.

## I. INTRODUCTION

The advent of functional magnetic resonance imaging (fMRI) revolutionized systems neuroscience in both humans and non-human primates (NHPs) [1], [2]. However, despite major advances in hardware, acquisition sequences, and analyses, efficiently measuring functional signals at the spatial scale required for circuit-level interrogation remains challenging. Functional ultrasound imaging (fUS) provides a solution by measuring at high spatiotemporal resolution (~150 x 150 x 100 µm3, ~2 Hz) cerebral blood volume (CBV) as a proxy for neural activity [3], with signals closely correlated with neuronal firing rates [4]. The technique has been successfully applied in mice [3], rats [5], ferrets [6], pigeons [7], marmosets [8], macaques [9], and humans [10]. Recent technical advances have enabled volumetric (3D) imaging (solely) in rodents [11]. Extending 3D-fUS to non-human primates would permit simultaneous 3D imaging of mesoscale cortical networks, reducing experimental time and enabling investigation of interactions among laminar and columnar scale modules. 3D-fUS can be further strengthened by adopting well-standardized fMRI practices in data processing and signal interpretation. A critical component of this process is the development of a hemodynamic response function for fUS (fUS-HRF).

The fMRI-HRF was created to relate the blood-oxygenation-level-dependent (BOLD) signal in fMRI to the underlying neural activity [12]. While imperfect, the HRF provided greater confidence in interpreting fMRI signals. However, the general applicability of the HRF is debated. Its generic use assumes consistency in the hemodynamic response across subjects, brain regions, and stimulus types, as well as stationarity in time, assumptions that have been questioned [13]. Several studies have indeed reported variability in HRFs [13], [14], [15], variability that may partly arise from the “impure” nature of the BOLD signal, which reflects a complex non-linear interplay of CBV, cerebral blood flow, and oxygen metabolism, rather than a single physiological parameter [16].

In NHPs, particularly macaques, monocrystalline iron oxide nanoparticle (MION) contrast agents have been used [1] to increase the contrast-to-noise ratio of fMRI signals, providing a purer measure of CBV [17], the same parameter measured by fUS [18]. Studies of the HRF in both BOLD-and MION-fMRI have employed similar experimental paradigms. In Boynton et al.’s [19] study in humans, black-and-white checkerboards were presented with variable stimulus durations. Similarly, Leite et al. [17] presented black-and-white pinwheel checkerboards to macaques, using stimulus durations of 4 and 60 s.

The goal of the present study is to explore the potential and sensitivity of 3D-fUS in macaques, with a particular focus on performing an in-depth analysis of the hemodynamic response profile. To address the discrepancies reported in fMRI-based HRF estimations, we expanded on existing HRF paradigms to determine whether a generalized HRF could be extracted or whether variability prevents such a generalization. By systematically varying stimuli, cortical regions, and subjects, we demonstrate that it is possible to extract a canonical HRF suitable for volumetric fUS in the macaque brain. This generalizable HRF facilitates the decoding of stimuli using more complex experimental designs and, importantly, enables more robust and reliable statistical analysis, much like the canonical HRF used in many fMRI studies.

## II. Methods

### A. Subject Details and Surgical Procedures

This study used three male rhesus macaques (*Macaca mulatta*, 7.5-9.5 kg, 11-15 years old). NHPs were group-housed (minimum cage size of 16-32 m^3^) with cage enrichments (wood, ropes, toys, foraging devices etc.) at the primate facility of the KU Leuven Medical School. Housing and handling were in accordance with the Weatherall report. NHPs were exposed to natural light and monitored artificial light for 12□hours every day. NHPs had unrestricted access to food and daily access to restricted volumes of fruits and water; during experiments NHPs had unlimited access to fluids as part of the performance-based protocol. Animal care and experimental procedures followed national and European regulations (Directive 2010/63/EU) and were approved by the Ethical Committee of KU Leuven. The health of the three subjects was monitored and maintained daily by trained technical staff, veterinary staff, and experimenters; all trained to work with NHPs. For each subject, MR-compatible headposts and recording chambers were surgically implanted and secured with ceramic screws and dental cement as previously described in Vanduffel et al., [1].

### B. Chamber Localizations

To optimize the position of the custom recording chamber (2.7 x 2.7 cm, inner diameter) for fUS experiments, we acquired anatomical T1-weighted MRI scans for each subject using a 3T horizontal-bore full-body scanner (Siemens, PrismaFit, Erlangen, Germany). Subjects were anesthetized with an intramuscular injection of 0.5 cc Domitor (1 mg/mL) and 0.25 cc Nimatek (100 mg/mL), followed by half this dose every subsequent hour. A custom-built 10-channel receiver coil was positioned around the subject’s head for scanning. Scans were acquired with the following parameters: 0.4 mm isotropic voxels, repetition time: 2700 ms, and echo time: 2.5 ms. To enhance the signal-to-noise ratio, ten scans of the brain were averaged. All measurements and subsequent analyses were conducted using 3D Slicer [20]. The averaged image was aligned to stereotaxic space using anatomical landmarks (i.e., the meatus acusticus and cribriform plate). The chamber was implanted covering the caudal parts of the intraparietal sulcus (IPS), lunate sulcus (LS), and superior temporal sulcus (STS), 24° from vertical in the coronal plane and 20° from vertical in the sagittal plane. A computed tomography (CT) scan was then performed using a Somatom Force Siemens CT scanner to verify the placement. CT parameters included a slice thickness of 0.4 mm, collimation of 64 × 0.6 mm, pitch of 0.85, and reconstruction using both bone and soft-tissue kernels. The CT scan was aligned to the anatomical MRI using the Advanced Normalization Tools (ANTs) registration module in 3D Slicer [21].

### C. Behavioral Paradigms

Subjects were trained on passive fixation tasks via operant conditioning, using liquid rewards (juice droplets). Visual stimuli were presented on a 32-inch screen 40 cm in front of the subject and aligned with their gaze. A virtual fixation window of 2° × 2° was defined at the center of the screen, and subjects were trained to maintain their gaze within this window for sustained periods of fixation as well as to rest their hands inside a box placed directly beneath and in front of their head [22], receiving juice rewards (contingent on correct fixation (gaze and hands)) in intervals between 800 and 1200 ms. All sessions were conducted with the subject in a head-fixed, sphinx position. A photocell, not visible to the monkeys, attached to the bottom-right corner of the monitor, signaled stimulus onset time and was additionally sent as a digital TTL pulse to our fUS acquisition system synchronizing behavioral and neural data. Eye tracking was performed at 120 Hz using an infrared-based system (ISCAN, Woburn, MA, USA).

For the variable duration experiment (Type I/II), subjects were required to maintain fixation on a central white dot against a gray background. At regular intervals, a circular stimulus (30° radius) appeared at the center of the screen for one of five durations: 2, 4, 8, 16, or 32 s. To allow full recovery of the hemodynamic signal following each stimulus, we used long inter-trial intervals (ITIs), ranging from 30 to 36.6 s in 0.83 s increments, with a probability weighting such that 30 s ITIs occurred nine times more frequently than 36.6 s ITIs, with intermediate durations linearly scaled in probability. Stimulus durations and inter-trial interval durations were randomized across trials. Runs were trial based and of random length (M1-M2-M3 mean: 13.69-8.18-8.61 min), with a random assignment of stimulus duration. Two days of data were collected per subject and only those runs with > 95 % (90% for M3) fixation accuracy were kept for further analysis. Only those trials in which the subject maintained fixation for the full trial were kept as “correct” trials for analysis. The minimum number of trials per stimulus duration, and not the total number of runs, are relevant for the analyses and are reported (n = 55 total, 11 per duration, per subject). The Type I stimulus was a colorful full-field radial checkerboard, presented as a wedge–ring pattern (48 wedge subdivisions over logarithmically spaced concentric rings), flashing (counter-phasing checkerboard flicker) at 10 frames per second for the specified stimulus duration. The Type II stimulus was one of two optic flow stimuli, either expanding or contracting dot motion, presented as black circular fields with moving white dots (dot radius = 0.25°, dot density = 20%), shown at 60 frames/s. Dot motion was defined relative to one of eight possible centers (at an eccentricity of ~ 9.4°) and followed a radial flow field. The speed of motion increased linearly with distance from the center of motion with a velocity gain of 0.5 degrees/s per degree of eccentricity.

A separate proof of concept version of the Type I task was run with single durations i.e., only 8 s stimuli were used. All other parameters are as described for the multi-duration experiment (except for the ITIs which ranged from 33 to 53 s with half-step intervals). This dataset was only recorded in M1. A single run of this data (15 trials) was used for further analysis (single run and single trial correlation). Correlation was defined as the normalized dot product between the voxel time series and a boxcar reference signal. Both the reference signal and voxel time-series were mean-centered and L2-normalized.

M1 performed an additional passive fixation task involving stimuli designed to map the horizontal and vertical meridian representations. Stimuli consisted of vertical and horizontal “bow tie” shapes reaching a 24.5° radius [1]. The patterns presented within the bow tie mask were identical to those used by Benson et al. [23] with each texture designed using a large variety of objects which are displayed at multiple scales on an achromatic pink-noise background. Objects were randomly selected and placed at random positions within the “bow tie” mask, and textures were updated at 4 Hz. Two temporal designs were used for this task: i) the single trial design: individual stimuli, either the vertical or horizontal bow ties, were shown for 4 s, followed by a 15.6 s intertrial interval, and ii) the four-trial design: four stimuli were shown in succession, each for 4 s with no inter-trial interval, in either the horizontal-vertical horizontal-vertical order, or the reverse. Each block of four stimuli was followed by a 15.6 s inter-trial interval. A total of 14 runs were acquired in a single session (7 per design).

### D. Image Acquisition

The imaging technique for 3D-fUS was adapted from [11]. The ultrasound probe was a matrix (32 rows by 32 columns) of piezoelectric elements, covering a surface area of 9.6 x 9.6 mm (Vermon France). The probe had a center frequency of 15 MHz, allowing penetration ~1 cm into the brain. A 1024-to-256 channel multiplexer selected an 8-row by 32-column rectangle and connected it to 256 transmit/receive channels (Vantage 256, Verasonics USA). The electronics were programmed to transmit a series of predefined plane waves at a fixed rate and to receive echoes from the brain. The echo data (referred to as RF data) were sent to a computer with 4 GPUs to reconstruct the vessel images in real time. For volumetric imaging, the plane wave emission was repeated multiple times, with the multiplexer aperture shifted by 4 rows with each emission (rows 1 to 8, 5 to 12, …, 25 to 32). Note that the apertures overlap. The received data were concatenated to obtain the RF signals for the full matrix, and the data from the overlapping rows were averaged. To increase the resolution, we used 7 plane waves at different angles (see Table 1) and combined the resulting images into a single composite image [11]. The composite image is acquired 200 times at a frequency of 500 Hz (~0.6 s acquisition time) to image the blood and brain tissue. The blood signal is extracted from the tissue using a singular value decomposition (SVD) filter. The intensity of the blood signal is then averaged to produce a microvascular image. These microvascular images are repeated every 0.6 s throughout the fUS acquisition.

**TABLE I.** Angles of the 7 plane waves (in Degrees)

|  |  |  |  |  |  |  |  |
| --- | --- | --- | --- | --- | --- | --- | --- |
| Angle X | -6 | -3 | 0 | 3 | 6 | 0 | 0 |
| Angle Y | 0 | 0 | 0 | 0 | 0 | -6 | 6 |

### E. High resolution vascular image and Image Registration

A high-resolution image was obtained using a 3D extension of the SUSHI method [24]. In this method, the intensity image is computed for each Doppler spectrum sub-band (n =20). Since the Doppler frequency is linked to blood velocity, the image of each sub-band shows vessels within a specific velocity range and is sparser than the original micro-Doppler image. For each sub-band image, we computed the vessel positions that optimize the experimental images with a sparse hypothesis. This optimization needs a point spread function (PSF) to reconstruct the image from the vessel positions. In our case, we used a gaussian function:

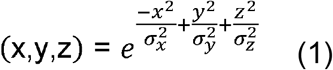

where *σ*_*x*_, *σ*_*y*_, *σ*_*z*_ are 180, 180 and 130 µm respectively.

To acquire the Doppler spectrum, we used the same procedure as in the functional acquisition, but with 11 angles instead of 7, and the frame rate was reduced to 1 Hz. The spectrum was averaged 100 times to improve the signal-to-noise ratio (SNR). The total acquisition time was 1 minute. Due to the inverse contrast between MRI and fUS modalities, where sulci appear dark in T1 images and bright in fUS (corresponding to high relative blood volume), manual alignment was possible through linear translations and rotations, without the need for non-linear warping. Final registration was completed using 3D Slicer (Fig 1B).

**Fig. 1.**
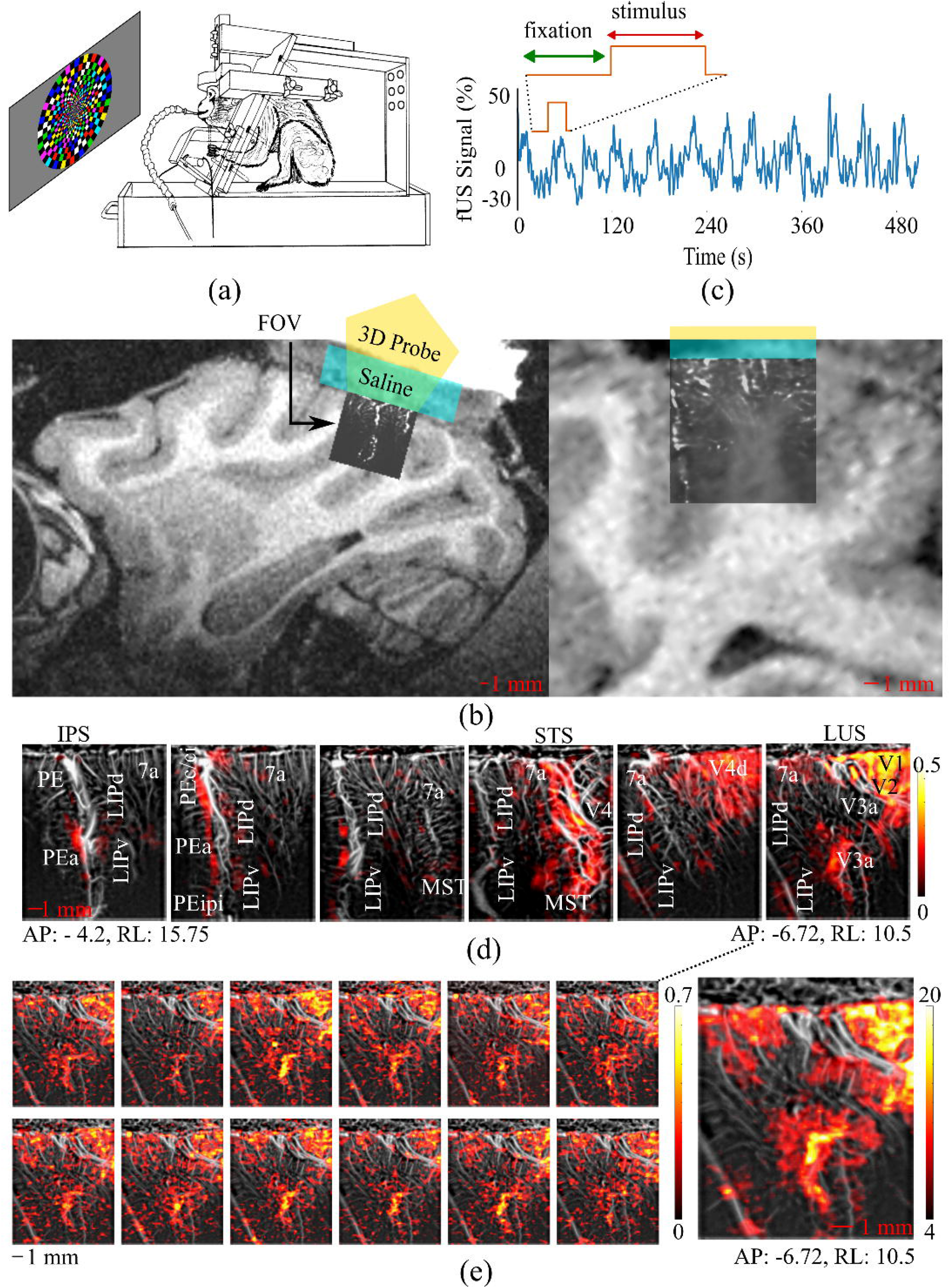
Experimental setup and 3D-fUS results. (a) Passive fixation task with variable-duration visual stimuli (2–32 s) and semi-randomized ITIs (33–36.6 s); reward contingent on eye fixation and hand position. (b) Alignment of the fUS FOV with sagittal MRI slice; chamber filled with saline to increase conductivity. (c) Single-voxel time course (area V2) showing percent signal change during 8-s checkerboard stimulation. (d) Coronal slices from a high-resolution microvascular fUS image with anatomical ROIs (PE, PEa, PEc/ci, PEipi, 7a, LIPd/v, MST, V4d, V3a, V1, V2) and functional correlation overlays (red/yellow) from a single run. (e) Single-trial correlation maps (15 trials) for the single-duration checkerboard experiment (left) and corresponding averaged z-score map (right).

### F. Between Run/Day Motion Correction

For each acquisition, we assessed potential movement of the probe within-day and between-days via the Python library for Advanced Normalization Tools (ANTS) [21] using the rigid registration method between the mean of the first run to each successive run. In the case where there was < 2 voxels of movement in any of the three translational axes within one day, no registration was applied; this was the case for M1 & M2. For all three subjects, between-day registration matrices were applied using linear interpolation. In the case of M3, as both within-day and between-day registration was necessary, one registration matrix was created considering the mean of the first run of the first day as the fixed position, and registering each subsequent run to this image, using a rigid registration with linear interpolation.

### G. Data Processing and Analysis (Variable Duration Experiment)

We acquired six sessions of the variable duration experiment, two per subject. Data were analyzed using custom Python scripts as follows. For each run, the Doppler signal was aligned with the timestamps (onset) of the corresponding stimuli. All runs from a single session were then concatenated into a continuous signal for analysis. fUS data were acquired at a stable rate of 1.67 Hz (M1) and 1.33 Hz (M2 and M3). Therefore, all data were first interpolated to 1.67 Hz. The signal was then segmented by stimulus duration, and trials were subsampled to ensure an equal number of trials per stimulus length (n = 11; total n = 55 for checkerboard, n = 55 for optic flow). Average baseline activity was defined as the mean activity measured during the final 15 s of the ITI following each 2 s trial, a period during which no stimulus-evoked activity was present. For each trial, the percent signal change was computed by comparing the trial signal to this baseline. To assess stimulus-driven activity, a per-voxel Wilcoxon signed-rank test was performed, comparing responses for each stimulus condition against the baseline, using a time window from 1.8 to 9 s post-stimulus onset, depending on the stimulus duration. This interval was chosen to capture the initial rise of the hemodynamic response. Effect size was estimated by converting the Wilcoxon W-statistic to signed z-scores.

All regions of interest (ROIs) were defined by manual selection of a rectangular region determined to be within the cortical ribbon of that anatomical area (as determined by CHARM) by the pattern of vascularization (small vessels perpendicular to the cortical surface). A ROI in M1’s visual area V2 was used to assess temporal differences between the neural responses evoked by separate stimuli, but within the same ROI, as it was universally activated across stimulus durations and types. A control ROI was placed in a portion of area V4d, which was non-responsive across stimulus durations and types, likely because it represents eccentricities beyond the range we tested. Within each ROI, average response profiles and standard error of the mean (SEM) were computed from the mean of all (significantly activated V2) voxels within a 0.018 mm^3^ volume across trials (Fig 2B/C). This analysis was performed in M1 (Fig 2) and M2 (not shown).

**Fig. 2.**
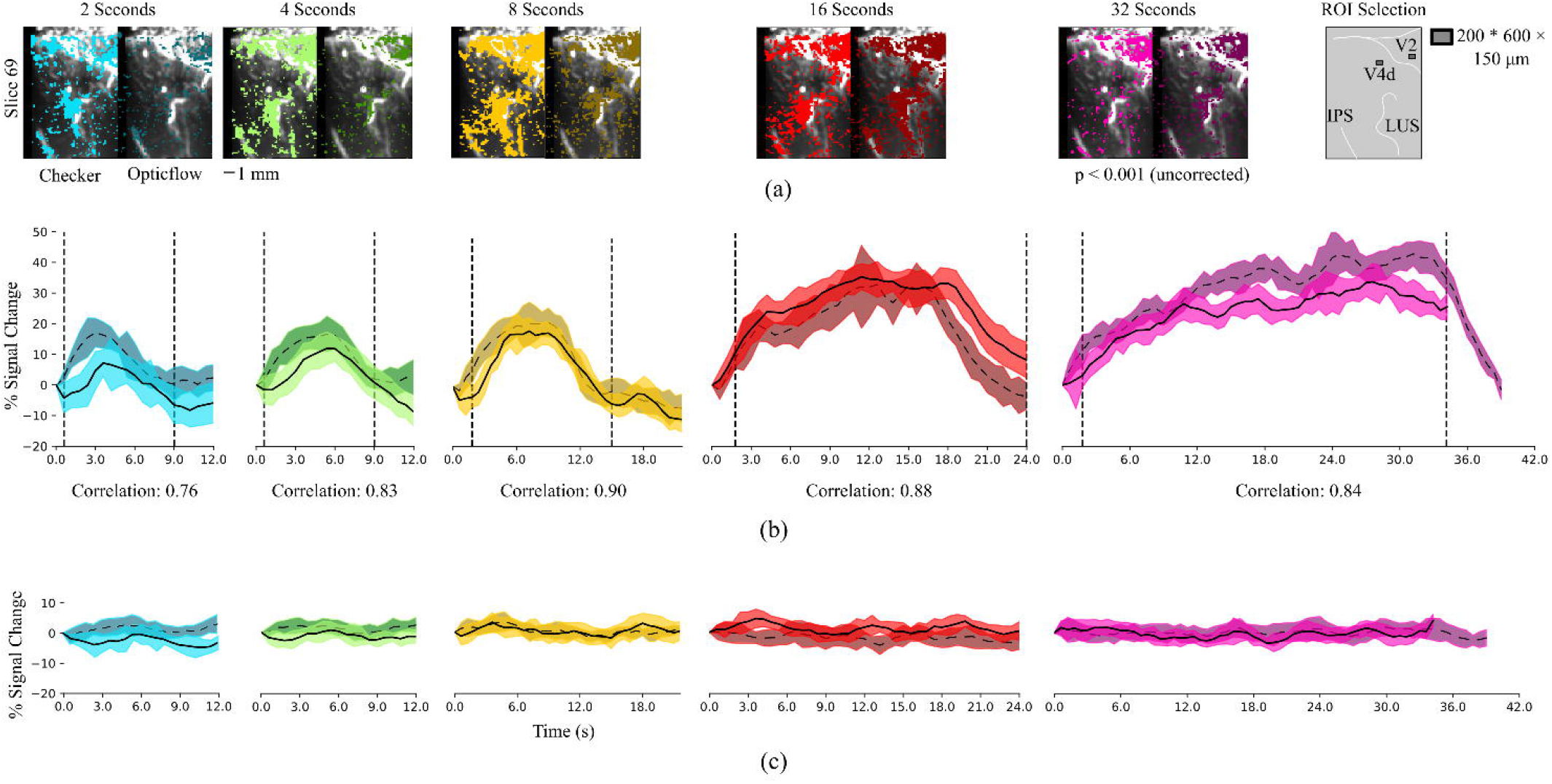
Stimulus-evoked fUS responses across durations and stimulus types. (a) Voxelwise Wilcoxon signed-rank test results (p < 0.001, uncorrected) for checkerboard (light) and optic-flow (dark) stimuli across durations (2–32 s), shown on a representative coronal slice. (b) Smoothed mean PSC (± SEM) from a V2 ROI (200 × 600 × 150 μm^3^) for each duration and stimulus type (checkerboard, solid; optic flow, dashed); Pearson correlations shown below. (c) Smoothed mean PSC (± SEM) from a control ROI in area V4d.

ROIs for further analysis were selected based on both anatomical location and functional activation (i.e., statistically significant voxels as determined by the Wilcoxon signed-rank test). For M1, ROIs were placed in primary visual cortex (area V1) and the medial superior temporal area (MST), and for M2, in posterior intra-parietal area (PIP), and lateral intraparietal area dorsal (LIPd). Within each ROI, average response profiles were computed from the mean of all significantly activated voxels within a 0.0945 mm^3^ volume across trials (Fig 3, Response Column).

**Fig. 3.**
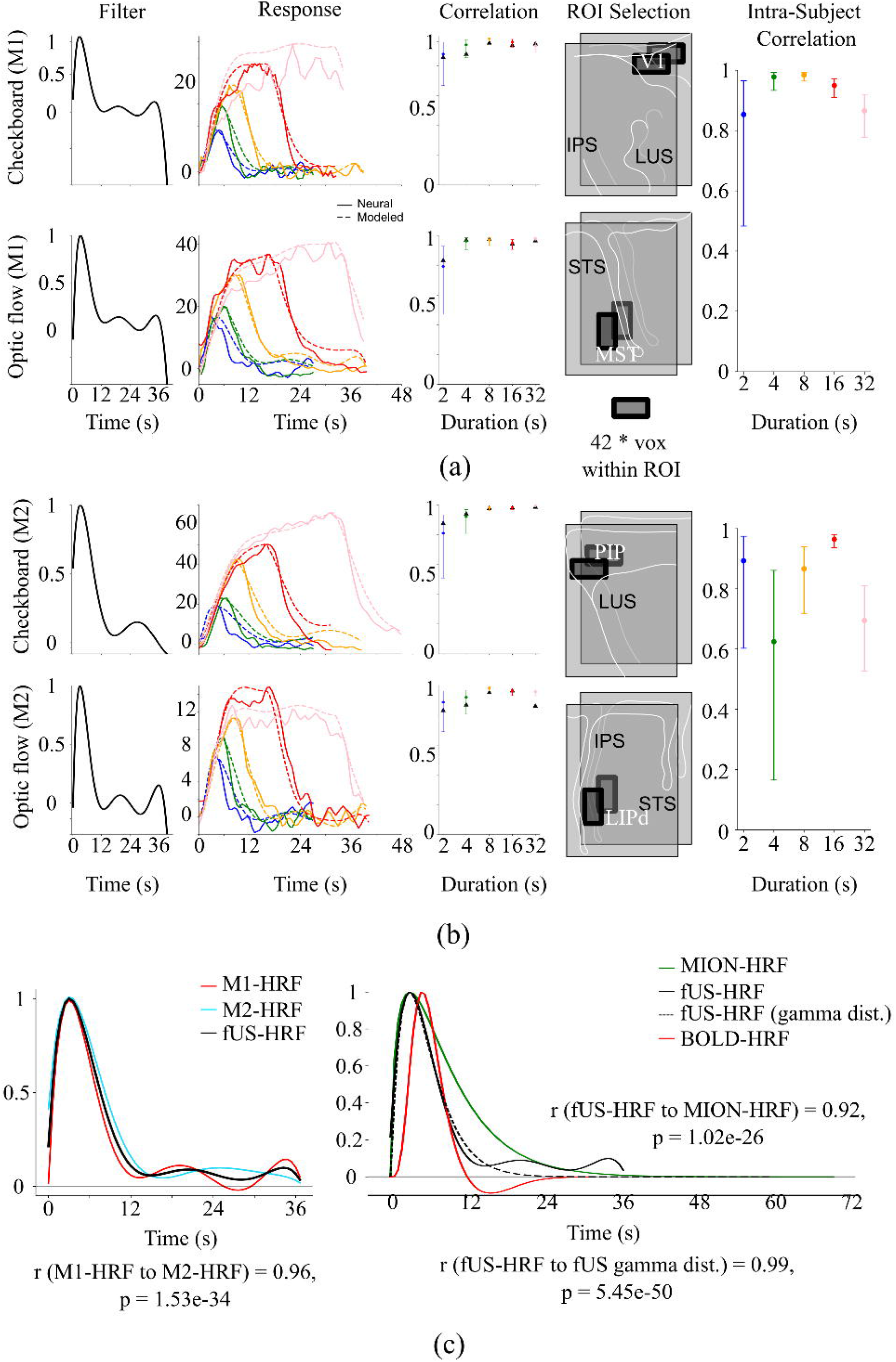
Estimation and validation of the macaque fUS-HRF. (a) Subject M1: exponential basis-function filters for checkerboard (V1) and optic-flow (MST) stimuli (Filter), simulated responses obtained by convolution with stimulus boxcars (Response), and Pearson correlations between simulated and real responses across durations (Correlation). (b) Same analyses as in (a) for subject M2 using ROIs in PIP (checkerboard) and LIPd (optic flow); black triangles indicate correlations obtained using the canonical fUS-HRF. (c) Subject-specific proto-HRFs (M1, blue; M2, red) and canonical fUS-HRF (black) compared with MION-fMRI (green) and BOLD-fMRI (red) HRFs; fUS-HRF gamma-function (black-dotted).

We estimated the fUS-HRF following the methods described in Leite et al. [17] with the addition of an optimization step based on a separate dataset. For each subject (M1 & M2), mean ROI responses representing the checkerboard (V1 & PIP) and optic flow (MST & LIPd) stimuli across five stimulus durations (2, 4, 8, 16, and 32 s) were used to estimate our fUS-HRF. The fUS-HRF was modeled as a linear combination of decaying exponential basis functions:

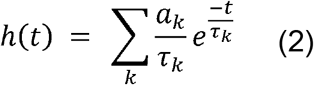

where *τ*_*k*_ are fixed time constants (in seconds) and *a*_*k*_ are amplitude coefficients estimated from data. The fUS-HRF was evaluated over a 40 s window at a temporal resolution of 0.6 s.

To fit this model to the neural responses, we derived the corresponding step responses for each exponential component. For a stimulus of duration d, the predicted response (i.e., after convolution) of a single exponential component is:

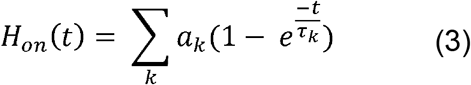

following stimulus onset, and

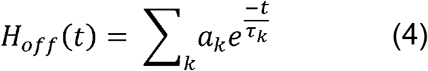

following stimulus offset.

For each stimulus duration, a design matrix was constructed containing: (1) one column per exponential component (i.e., the step response evaluated at each [Equation]); (2) an intercept term; and (3) a linear trend regressor to capture slow drifts. For each subject/stimulus pair, the design matrices from all five stimulus durations were vertically concatenated, and the corresponding empirical responses were concatenated into a single response vector. The amplitude parameters [Equation] were estimated using ordinary least squares (OLS). This procedure yielded one HRF estimate per subject/stimulus pair (four total) (Fig 4, Filter Column). The final fUS-HRF used for subsequent analyses was computed as the average of these four subject/stimulus-specific HRFs (Fig 4C).

**Fig. 4.**
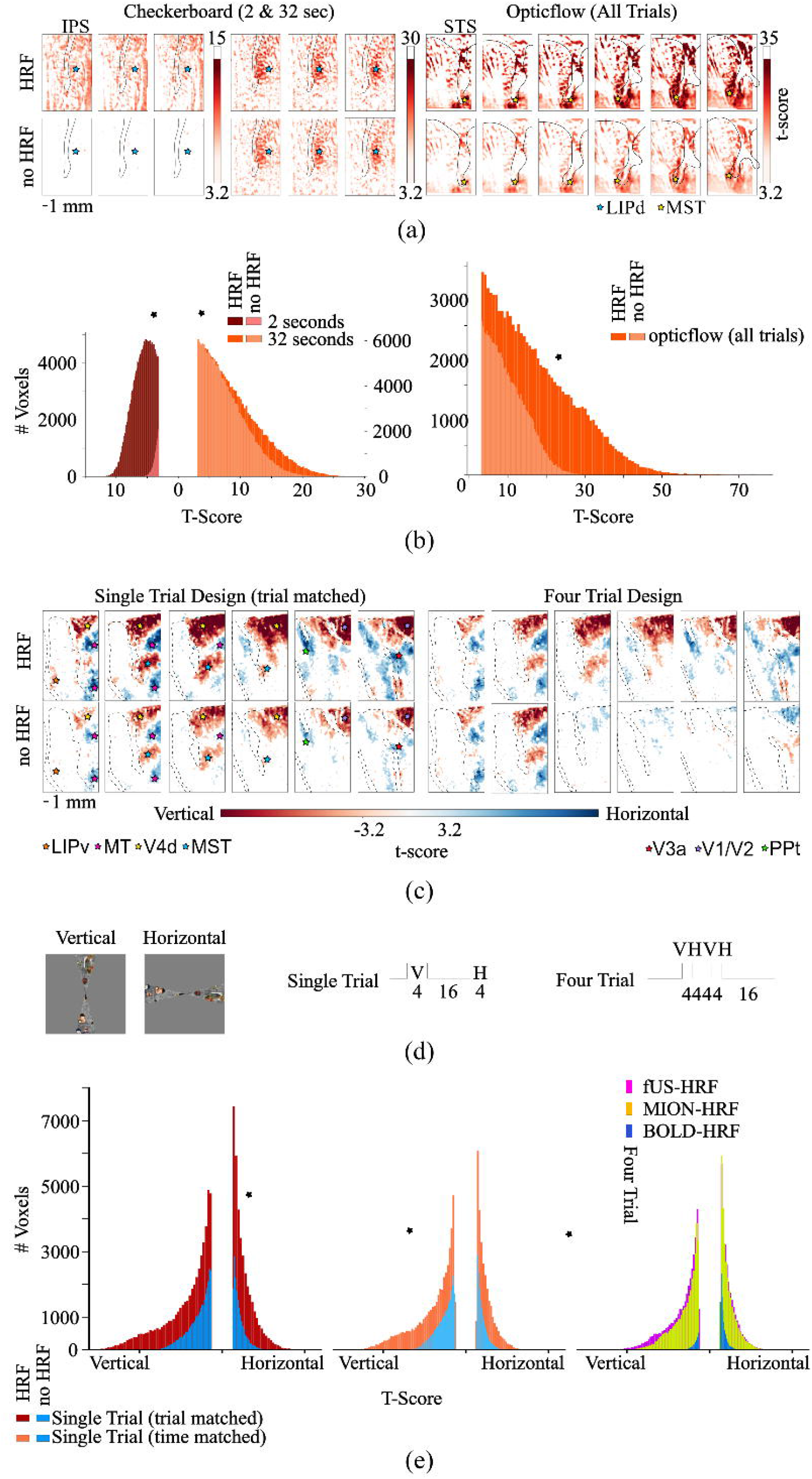
Effect of fUS-HRF on GLM analyses. (a) GLM t-score maps from subject M3 for checkerboard (2-s and 32-s) and optic-flow stimuli analyzed with (top) and without (bottom) HRF convolution (p < 0.001, uncorrected); selected coronal slices shown (14, 15, 16 and 54, 55, 56, 57, 58); stars indicate anatomical ROIs (LIPd: blue, MST: yellow). (b) Distributions (bin size = 100) of significant positive t-scores corresponding to (a); paired t-tests compare HRF and no-HRF conditions (p < 0.001). (c) GLM t-score maps from M1 for meridian-mapping tasks using single-trial and four-trial temporal designs, analyzed with and without HRF convolution (slices: 40, 45, 50, 55, 60, 65), p < 0.001, uncorrected). Stars indicate relative position of different anatomical areas, (LIPv: orange, V4d: yellow, PPT: green, V1/V2: purple, MT: pink, MST: blue, V3a: red) (d) Meridian-mapping task designs. (e) Distributions (bin size = 100) of significant positive t-scores for single-trial (darker color indicates HRF usage) and four-trial designs (fUS-HRF: pink, MION-HRF: yellow, and BOLD-HRF: blue; paired t-test: p < 0.001, indicated by a black star).

For the first pass in constructing the fUS-HRF, a set of τ values representing equidistant time steps across the full plausible neural response window (0 to 42 s) was input. To optimize and evaluate candidate fUS-HRFs, we implemented an optimization procedure based on a general linear model (GLM) analysis of a separate meridian mapping dataset (described above). The objective was to maximize the number of suprathreshold voxels achieved in the dataset irrespective of sign (suprathreshold was defined as p < 0.001). Optimization was performed using a stochastic local search strategy over 1000 iterations. At each iteration, candidate time constants were generated by adding Gaussian perturbations (standard deviation = 3 s) to the current best solution. The time constants were constrained to the interval 0.6 - 90 seconds and sorted in ascending order. For each candidate parameter set, the fUS-HRF was re-estimated from the empirical step responses as described above, and the GLM was recomputed.

### H. General Linear Model

In brief, the GLM was implemented in Python following standard first-level fMRI analysis procedures, using an OLS approach. This method was based on the framework of Statistical Parametric Mapping, specifically [25]. A design matrix was constructed, including the main regressor of interest (task events/duration convolved with our fUS-HRF), nuisance regressors (i.e., pupil size and eye position), and a constant term. For each voxel, regression coefficients were estimated directly from the voxel’s timeseries using OLS. Residuals were computed, and error variance was estimated from the unexplained signal. T-statistics and associated p-values were then derived for the regressor of interest.

As our primary goal was not to interpret the specific GLM results, but rather to evaluate the performance of our HRF in a new subject (and in a separate task), we conducted a GLM sensitivity analysis in subject M3 (Fig 4A/B) using the previously described duration tasks, as well as the meridian mapping task recorded in M1 (Fig 4C/D/E). To assess the impact of our HRF model, we performed these analyses both with HRF-convolved regressors and with un-convolved boxcar style regressors. Paired t-tests using the scipy stats module in python were used to assess whether there was a significant difference between the relevant t-score distributions.

For both types of the variable duration task recorded in M3, the signal from each run was concatenated, and a design matrix was created that included the event onsets and durations for each trial. Pupil size and the x, y positions of one eye were included as regressors of no interest. This basic GLM setup was applied both with and without HRF convolution. The resulting t-score maps were thresholded (p < 0.001, uncorrected) and masked for positive values to assess the spread of significant positive t-scores. To assess the effect of HRF usage on the analysis of stimuli of different lengths, for the checkerboard stimuli we applied this logic to an evenly matched subset of trials (n = 8) of the 2 s and 32 s durations (Fig 4A, left).

For both temporal designs of the meridian mapping task, as with the duration task, the GLM was applied to both paradigms (single-trial and four-trial variants of the meridian mapping task) with and without HRF convolution. A simple (horizontal versus vertical meridian) contrast was used. The resulting t-score maps were thresholded (p < 0.001, uncorrected) and the distribution of t-scores was calculated for each condition. Importantly, we performed these GLMs with either the number of trials matched (for the single trial versus four-trial design), or with the total acquisition time matched between the two designs (i.e., total number of 3D-fUS frames). For the comparison and evaluation of our fUS-HRF, we also analyzed this dataset (specifically the four trial designs) with the macaque BOLD-HRF and MION-HRF (Fig 4E, rightmost distribution).

## III. Results

### A. 3D-fUS in Macaques

We adapted key concepts from NHP-fMRI paradigms to achieve stable 3D-fUS recordings over extended periods (M1: > 22 months, M2: > 10 months, M3: > 6 months). A custom NHP chair [1] provided secure head fixation, while a juice reward system delivered controlled reinforcements during operant behavioral tasks. Juice delivery was contingent on sustained fixation (gaze/hands) [22], which minimized motion-related artifacts during fUS acquisition (Fig 1A). For fUS recordings, a custom chamber was implanted over the posterior sectors of the IPS and STS, with an adaptor designed to securely hold the fUS probe in place (Fig 1B). In all experiments, we imaged a 3D volume of approximately 1 cm^3^ in the macaque brain (100 x 150 x 150 µm^3^ voxel size) at a frame rate of ~1.67 Hz, while the monkeys performed passive fixation tasks (see Methods). This setup allowed for daily probe placement at a consistent position. Within-session movement were limited to 2 voxels, and across sessions, we observed a maximum shift of 6 voxels along the medio-lateral axis. Image registration was applied across all runs acquired over the two recording days. To quantify positional variation of the probe, we assessed these metrics (see Methods: Between Run/Day Motion Correction) in an independent multi-day experiment (minimum 4 days). Positional variation expressed in mm^3^/degrees, (translation/rotation), was M1: 2.745e^-3^/5.625 e^-4^, M2: 4.1625e^-3^/1.98e^-3^, M3: 7.5825e^-3^/3.0375e^-3^.

To register the fUS images with an anatomical atlas, we acquired a high-resolution microvascular image achieving substantially better contrast for identifying the cortical ribbon. The vasculature features in this image facilitated delineation of the pial surface and the grey-white matter boundary, which is crucial for accurate registration of the vascular maps to a T1-weighted anatomical MR image (Fig 1B). We successfully registered the CHARM atlas [26], [27] to the anatomical fUS images (Fig 1D).

When a visual stimulus (i.e. a colorful checkerboard, 8 s duration) was presented, the fUS signal in early visual cortex increased by approximately 25 to 50%, as evident in the single voxel time course in area V2 (Fig 1C). Fig 1D shows correlation maps (see Methods) from a single run of the single duration visual task on selected coronal slices (high-resolution microvascular images). In addition to pronounced fUS responses in early visual areas such as V1 and V2, the checkerboard stimulus also induced spatially specific activity in higher-order areas, including medial superior temporal area (MST) and various superior parietal (PE) sectors. Single-trial correlation maps (n =15) revealed correlation coefficients as high as r = 0.7 in the most active regions (V1/V2), highlighting the excellent reproducibility of the fUS results (compare the individual trial maps in the left panel of Fig 1E). Consistent with this, the average z-score across single trial maps reached values up to 40, with most highly activated regions exhibiting z-scores around 20.

### B. Characterizing the fUS-HRF

To derive an HRF applicable across subjects, brain areas, and stimulus types, we recorded fUS signals from two macaques. NHPs were presented with two types of visual stimuli — flashing checkerboards and optic flow patterns, each displayed for multiple durations. Stimulus durations were randomized, and ITIs (i.e., the interval between stimulus offset and stimulus onset of the next trial) were randomized in the range from 30 to 36.6 s.

To assess the intra-subject and intra-areal reproducibility of fUS responses, we used an equal number of trials (n = 11 per stimulus type and duration) and compared the spatio-temporal fUS responses between the visual and the baseline trials (no stimulus). Both checkerboard and optic flow stimuli, across stimulus durations, activated similar regions in early and extrastriate visual cortex, including V4, MT and MST, although to varying extents, (see selected slice in Fig 2A). Statistical comparisons were performed using Wilcoxon signed-rank tests with FDR correction, using a threshold of p <0.001 (uncorrected) for individual durations.

We extracted time courses (percent signal change (PSC)) for each stimulus type and duration from a small V2 ROI (200 × 600 × 150 μm^3^; Fig 2B; see its location in the right panel of Fig 2A). Pearson correlations between responses evoked by the checkerboard and optic flow stimuli across durations ranged from 0.76 to 0.9, indicating strong similarity in response dynamics between the two stimuli. These results show temporally consistent responses to different visual stimuli within the same subject and ROI. Time courses extracted from an equally sized control ROI of area V4d showed no stimulus-evoked fUS activity (Fig 2C; see its location in the right panel of Fig 2A).

To explore inter-subject and inter-ROI responses to different stimuli, we calculated Pearson correlations between the time courses evoked by checkerboard and optic flow stimuli for matched stimulus durations within individual subjects (N = 2) across different cortical areas. In both regions (V1 and MST, M1), fUS response amplitudes increased monotonically with stimulus duration, from approximately 5% signal change for the shortest duration to > 30% for the longest. Additionally, responses in V1 to checkerboard stimuli and in MST to optic flow stimuli were highly correlated, with Pearson correlation coefficients exceeding 0.80 in all cases (p < 0.001; Fig 3A, intra-subject correlations).We repeated the same analysis in M2, using area posterior intraparietaal area (PIP) for the checkerboard stimulus and dorsal lateral intraparietal area (LIPd) for the optic flow stimulus (Fig 3B). While the amplitude of the PSC differed between regions, the temporal response patterns were qualitatively similar. This similarity was supported by high Pearson correlations exceeding 0.6 (p < 0.001). Replicating these results across two subjects suggests that distinct brain areas exhibit comparable temporal response patterns, even when driven by different visual stimuli.

Using the time courses from the analyses above, we derived an HRF intended to generalize across stimuli, regions, and subjects. For each stimulus-subject pair, we computed an exponential basis function (see Methods), yielding four proto-HRFs (Fig 3, first “Filter” column). We then averaged across stimuli within each subject to generate subject-specific HRFs, which were highly similar between the two animals (r = 0.96, p = 1.53e-34; Fig 3C, left).

Next, we tested the subject-specific HRFs by calculating Pearson’s correlations between the simulated responses (stimulus paradigm convolved with the HRF) and the actual response for each stimulus, duration, and subject. Correlations were typically > 0.8 across subjects (Fig 3, Correlation Column), though they varied by duration. Importantly, the shape of the response, including the rising slope, peak, plateau (if present), and decay slope were similar for both functions. These were then averaged to create a canonical macaque fUS-HRF. This fUS-HRF can be approximated with a gamma distribution probability density function defined as:

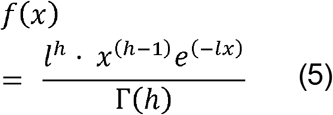

with h = 2.1 and l = 2.7 (Fig 3C, right). The correlation coefficient between the fitted curve and the experimental HRF is 0.99 (p = 5.45 e-50). Notice that the gamma distribution probability function is also used to fit the HRF for MION-fMRI, but the parameters are different (h = 1.55, l= 0.0127) and the generated curve is longer (Fig 3C, right). Even so, the correlation between the fUS-HRF and MION-HRF is quite high (r = 0.92, p = 1.02e-26), and the time to peak is identical. In contrast, the macaque BOLD-HRF presents a significantly delayed time to peak, as well as a faster decay, achieving a considerably lower correlation with the fUS-HRF (r = 0.83, p =1.32e-14).

As a reverse validation of the “canonical” fUS-HRF, we convolved a boxcar stimulus for each stimulus duration and compared the result to the true neural response for each monkey, stimulus type, and duration (Fig 3, Correlation column, black triangles). In M1, correlations between the canonical fUS-HRF and the true neural responses were consistently above 0.8, with most values within 5% of those obtained using the M1-specific HRF. In four conditions (checkerboard 32 s, optic flow 2, 4, and 8 s), the canonical fUS-HRF yielded higher correlations than the M1-specific HRF. A similar pattern was observed in M2: all correlations exceeded 0.8, and in two conditions (checkerboard 2 and 4 s), the canonical fUS-HRF yielded higher correlations than the M2-specific HRF. These results show that the HRF derived from data across two monkeys, four areas, and two stimuli, generalizes well.

### C. Application in general linear models

A well-fitted HRF should improve the temporal alignment of the actual brain response and the stimulus paradigm, increasing the statistical power of GLM analyses. To test this, we first performed GLM analyses both with and without fUS-HRF convolution, using data from a subject (M3) that did not contribute to the creation of the fUS-HRF (Fig 4A). Separate analysis strategies were applied for the two stimulus types. For the checkerboard stimuli, we selected equal numbers of trials with 2 s and 32 s durations and applied our HRF and no-HRF GLM pipeline. Fig 4a (left) shows t-score maps from the resulting GLMs on representative anatomical slices (14-16). Although both stimulus durations activated similar regions of the cortical ribbon (specifically LIPd), CBV responses to the short-duration stimuli were largely undetectable without HRF convolution. This discrepancy was reduced for longer block-style stimuli (32 s). For the optic flow stimuli, we pooled all stimulus durations for both GLMs. Here as well, omission of the HRF resulted in reduced t-score (slices 54-59), reflected as a desaturation of the statistical maps. Consistent with these voxel-based observations, the distributions of t-scores across the entire fUS block differed profoundly between the HRF and the no-HRF based analyses (Fig 4B). In all cases, the use of the fUS-HRF led to statistically significant differences (paired t-test) in the t-score distributions (checkerboard 2 s: t-value= 438.34, pvalue = 0.0, df = 29299; checkerboard 32 s: t-value =376.21, pvalue = 0.0, df = 166047; optic flow: t-value: 450.81, pvalue = 0.0, df = 91795).

We analyzed data from a different task in M1 that used distinct stimuli and a completely different temporal structure. Specifically, horizontal and vertical wedges were alternated using two temporal designs. GLM results for both designs, using matched numbers of trials and performed with and without fUS-HRF convolution, are shown in Fig 4C. In both single-and four trials designs, the use of the fUS-HRF led to increased statistical power (Fig 4C, E), corroborating the findings from M3’s checkerboard and optic flow experiments with varying stimulus durations. Specifically, both t-scores and the number of significant voxels increased when the fUS-HRF was included. Notably, in the four-trial design, many stimulus-driven activations were either missed or incorrectly identified when the fUS-HRF was omitted, underscoring the critical role of the HRF in this design. Note that such designs, which lack consistent ITIs, are commonly used in functional imaging studies due to their greater time efficiency.

We performed the same analysis on the four-trial design using the MION-and BOLD-HRFs (Fig 4E, rightmost distribution), which highlight two key findings. Using our fUS-HRF, we obtained a significantly better distribution of t-scores than with the MION-HRF despite the similarity in their shapes (paired t-test (over absolute t-score values): t-value= 161.383, p-value=0.0, df=61692). The BOLD-HRF, while producing higher t-scores than a simple boxcar (not shown), performed significantly worse in a paired t-test than both the fUS-or MION-HRFs (fUS-BOLD: t-value=96.1, pvalue=0.0, df=9096, MION-BOLD: t-value=98.5, pvalue=0.0, df=9097).

## IV. Discussion

In this study, we present the first successful implementation of volumetric (3D) fUS in primates by demonstrating highly consistent, reproducible imaging results in three subjects. We detected reliable 3D-fUS signals even for single stimulus presentations, highlighting the exceptional sensitivity of 3D-fUS. The 2 x 2 cm^2^ craniotomy required for this study remained stable for over 22 months, with no observable changes in the brain images, functional responses, or animal health. In addition, we produced high-resolution vascular images that enable precise anatomical identification and facilitate registration with a standard T1 anatomical MR image and anatomical atlases. Concerning the functional responses obtained with 3D-fUS, we observed signal increases of up to 50% for long visual stimulus and ~20% for shorter stimuli, which were readily detected in single trials. When functional activity was averaged across multiple trials, the signal proved to be highly stable, with a signal-to-noise ratio of approximately 30% for single voxels within an active region. We observed temporally consistent responses across stimuli within individual subjects (Fig 2), across cortical areas within a subject (Fig 3), and across subjects (Fig 3). Our within-subject HRFs exhibited remarkably similar shapes across two animals and across the two visual stimuli tested. Comparisons between the measured signal and predictions generated using within-subject HRFs yielded consistently high correlations (>0.8), while the correlation between the two proto-HRFs was close to 1.

One of the primary applications of the HRF is within GLMs, where it is used to generate a predicted temporal response. We show that the macaque fUS-HRF is significant and necessary to GLM analyses in three ways. First, we show that the HRF generalizes to a subject that did not contribute to its creation. This is critical, as a common criticism of fMRI-HRFs is their limited applicability across subjects [13], [15]. Second, in event related designs, while spatial activation patterns appear qualitatively similar with and without HRF convolution, inclusion of the HRF leads to statistically significant increases in both the number of significant voxels and their associated t-scores, improving statistical sensitivity. For short-duration trials, inclusion of the HRF becomes essential, as stimulus-evoked activity cannot be reliably detected without it. Finally, we demonstrate the critical importance of using the fUS-HRF in more temporally complex stimulation paradigms. In event-related designs without ITIs, HRF convolution is required to disentangle overlapping responses to successive stimuli. These results show that the fUS-HRF not only enables more complex and flexible experimental paradigms but also provides greater statistical confidence in the interpretation of results.

Our estimated fUS-HRF is significantly shorter than the MION-HRF. This difference in duration, however, is primarily due to the prolonged decay phase that characterizes the MION-HRF [1], [17]. The time to peak is identical between the two signals (3 s) and the relative shape of the initial rise is closely matched. In contrast, the macaque BOLD-HRF has a significantly delayed rise (time to peak: 4.8 s) and a faster decay [2]. These distinctions highlight that both fUS and MION-fMRI primarily measure CBV, whereas the BOLD signal reflects a more complex interaction of cerebrovascular dynamics. Our data support the view that fUS provides a direct measurement of CBV, while also revealing subtle but meaningful differences from MION-based signals. One plausible explanation for these differences between the fUS-/MION-HRFs is that fUS is more sensitive to arterial than venous compartments. fUS detects CBV changes via the Doppler effect, which primarily captures blood moving faster than approximately 4 mm/s in the axial direction [28], preferentially reflecting arteriolar flow while largely excluding venous components. During brief neural activation, the initial hemodynamic response consists of arteriolar vasodilation, which is detected by both fUS and fMRI. Following stimulus offset, however, arteriolar constriction redistributes blood into the slower venous network. While this prolonged venous component remains detectable with contrast-agent enhanced fMRI, it is attenuated, or even absent, in fUS measurements, resulting in a shorter HRF. This selective sensitivity highlights fUS’s potential for capturing high-temporal-resolution CBV dynamics that may be blurred in contrast-agent enhanced fMRI.

The shorter duration of the fUS-HRF has practical implications for experimental designs. First, in an event-related design, faster return-to-baseline allows for shorter ITIs. Second, it facilitates the separation of overlapping responses in rapid event-related designs, or in block designs without blank stimulus intervals. Finally, in brain decoding applications, the reduced HRF duration increases the number of discriminable events per unit time, thereby improving the temporal resolution and decoding performance. The approximated gamma function of the fUS-HRF provides a practical and convenient tool for implementing the fUS-HRF in data processing and GLM analyses.

This study has three practical limitations. The typical imaged volume (~1 cm^3^) is sufficient to visualize a large portion of the cortex, but sub-cortical regions remain out of reach, and still, the FOV remains far smaller than the whole-brain coverage of fMRI. Moreover, while fMRI in macaques cannot be considered non-invasive, 3D-fUS requires a craniotomy, representing an additional experimental and health consideration. Moreover, despite our extensive approach to estimate the fUS-HRF, more definitive conclusions will require replication across a larger number of subjects, multi-modal stimuli, and measurement from additional brain areas. Despite these limitations, the establishment of a macaque-specific HRF for fUS provides a robust foundation for both hypothesis-driven analyses (e.g., GLMs) and data-driven decoding approaches, enhancing reproducibility and comparability across studies. By enabling high-resolution measurements of sub-millimeter functional domains, such as laminar differences and mesoscale circuits, 3D-fUS has the potential to become a uniquely powerful tool for advancing systems-level neuroscience in primates.

## Acknowledgment

The authors thank C. Fransen, I. Puttemans, A. Hermans, S. Verstraeten, S. Riyahi, W. Depuydt, and M. De Paep for technical and administrative support.

